# APOBEC3 degradation is the primary function of HIV-1 Vif for virus replication in the myeloid cell line THP-1

**DOI:** 10.1101/2023.03.28.534666

**Authors:** Terumasa Ikeda, Ryo Shimizu, Hesham Nasser, Michael A. Carpenter, Adam Z. Cheng, William L. Brown, Daniel Sauter, Reuben S. Harris

**Author notes:** ^#^ These authors contributed equally (Terumasa Ikeda, Ryo Shimizu).

## Abstract

HIV-1 must overcome multiple innate antiviral mechanisms to replicate in CD4^+^ T lymphocytes and macrophages. Previous studies have demonstrated that the APOBEC3 (A3) family of proteins (at least A3D, A3F, A3G, and stable A3H haplotypes) contribute to HIV-1 restriction in CD4^+^ T lymphocytes. Virus-encoded virion infectivity factor (Vif) counteracts this antiviral activity by degrading A3 enzymes allowing HIV-1 replication in infected cells. In addition to A3 proteins, Vif also targets other cellular proteins in CD4^+^ T lymphocytes, including PPP2R5 proteins. However, whether Vif primarily degrades only A3 proteins or has additional essential targets during viral replication is currently unknown. Herein, we describe the development and characterization of *A3F*-, *A3F/A3G*-, and *A3A*-to-*A3G*-null THP-1 cells. In comparison to Vif-proficient HIV-1, Vif-deficient viruses have substantially reduced infectivity in parental and *A3F*-null THP-1 cells, and a more modest decrease in infectivity in *A3F/A3G*-null cells. Remarkably, disruption of A3A–A3G protein expression completely restores the infectivity of Vif-deficient viruses in THP-1 cells. These results indicate that the primary function of Vif during HIV-1 replication in THP-1 cells is the targeting and degradation of A3 enzymes.

**Importance:** HIV-1 Vif neutralizes the HIV-1 restriction activity of A3 proteins. However, it is currently unclear whether Vif has additional essential cellular targets. To address this question, we disrupted *A3A* to *A3G* genes in the THP-1 myeloid cell line using CRISPR and compared the infectivity of wildtype HIV-1 and Vif mutants with the selective A3 neutralization activities. Our results demonstrate that the infectivity of Vif-deficient HIV-1 and the other Vif mutants is fully restored by ablating the expression of cellular A3A to A3G proteins. These results indicate that A3 proteins are the only essential target of Vif that is required for HIV-1 replication in THP-1 cells.

## Introduction

The apolipoprotein B mRNA editing enzyme polypeptide-like 3 (APOBEC3, A3) family of proteins comprise seven single-strand DNA cytosine deaminases (A3A–A3D and A3F– A3H) in humans (1-3). A3 enzymes have broad and essential roles in innate antiviral immunity against parasitic DNA-based elements (4-6). Retroviruses are sensitive to A3 enzyme activity due to the obligate step of reverse transcription during viral replication that produces single-stranded cDNA intermediates. These viral cDNA intermediates can act as substrates for A3 enzymes, as demonstrated by C-to-U deamination resulting in G-to-A mutations in the genomic strand. To date, the best-characterized substrate of A3 enzymes is human immunodeficiency virus type 1 (HIV-1). In CD4+ T lymphocytes, four A3 proteins (A3D, A3F, A3G, and stable A3H haplotypes) restrict HIV-1 replication by mutating viral cDNA intermediates and by physically blocking reverse transcription (7-14). A3 enzymes have a preference for specific dinucleotide motifs (5′-C<u>C</u> for A3G and 5′-T<u>C</u> for other A3 enzymes) at target cytosine bases, which appear as 5′-<u>A</u>G or 5′-<u>A</u>A mutations in the genomic strand (7, 8, 15, 16).

Virus-encoded virion infectivity factor (Vif) functions in disrupting the activity of A3 enzymes. Vif forms an E3 ubiquitin ligase complex that degrades A3 enzymes through a proteasome-mediated pathway (2, 3, 17, 18). The central domain of this complex is a Vif heterodimer with the transcription factor, CBF-β, which stabilizes Vif during disruption of A3 protein activity (19, 20). Vif also suppresses the transcription of A3 enzymes by hijacking RUNX/CBF-β complex (21). In addition to these Vif-dependent mechanisms, HIV-1 reverse transcriptase and protease have been shown to disrupt the activity of A3 enzymes via Vif-independent mechanisms (22, 23). Recently, functional proteomic analyses have demonstrated that Vif has several target proteins, including the PPP2R5 family of proteins, in CD4^+^ T cell lines and lymphocytes (24, 25). These findings indicate that Vif may have additional essential target proteins during HIV-1 replication in infected cells.

We previously reported that endogenous A3G protein contributes to HIV-1 restriction in a deaminase-dependent manner in THP-1 cells (26). Although disruption of the *A3G* gene nearly eliminates viral G-to-A mutations, Vif-deficient HIV-1 virions have 50% lower infectivity than wildtype HIV-1 or mutants selectively lacking A3G degradation activity (26). These results indicated that Vif-mediated inhibition of A3G and at least one additional A3 protein is required for efficient HIV-1 replication.

In the present study, we evaluate the effects of other A3 proteins on HIV-1 infectivity by developing and characterizing *A3F*-, *A3F/A3G*-, and *A3A*-to-*A3G*-null THP-1 cells using HIV-1 Vif mutants with selective A3 neutralization activities. In comparison to wildtype HIV-1, Vif-deficient HIV-1 infectivity is strongly inhibited in *A3F*-null THP-1 cells and modestly inhibited in *A3F/A3G*-null THP-1 cells. In contrast, an HIV-1 Vif mutant selectively lacking A3F degradation activity had comparable infectivity to wildtype HIV-1 in *A3F*-null THP-1 cells and 50% infectivity in parental THP-1 cells, indicating that A3F protein contributes to HIV-1 restriction in THP-1 cells. Furthermore, Vif-deficient HIV-1 infectivity is comparable to wildtype HIV-1 in *A3A*-to-*A3G*-null THP-1 cells. These results demonstrate that A3 proteins are the primary target of HIV-1 Vif during virus replication in THP-1 cells.

## Results

### Endogenous A3H is not involved in HIV-1 restriction in THP-1 cells

THP-1 cells express significant levels of *A3B*, *A3C*, *A3F*, *A3G,* and *A3H* mRNA (26). The results of our previous study indicated that A3G and at least one additional A3 protein are involved in HIV-1 restriction in THP-1 cells (26). Variations in the amino acid sequence of A3 family proteins are known to influence HIV-1 restriction activity (27), and the *A3H* gene is the most polymorphic of all human *A3* genes (10, 22, 28, 29). The A3H allele is grouped into stable and unstable haplotypes according to the combination of amino acid residues at positions 15, 18, 105, 121, and 178 (10, 22, 28, 29). Stable A3H haplotypes are active against HIV-1 whereas unstable A3H haplotypes have absent or minimal activity as they encode proteins with low stability (9, 10, 22, 29, 30). To determine *A3H* genotypes, we sequenced *A3H* cDNA from THP-1 cells. Sequencing data identified an unstable haplotype in the THP-1 genome, termed A3H hapI (**Fig. 1A**). These data suggest that endogenous A3H protein has minimal restriction activity against Vif-deficient HIV-1 in THP-1 cells.

**Figure 1.**
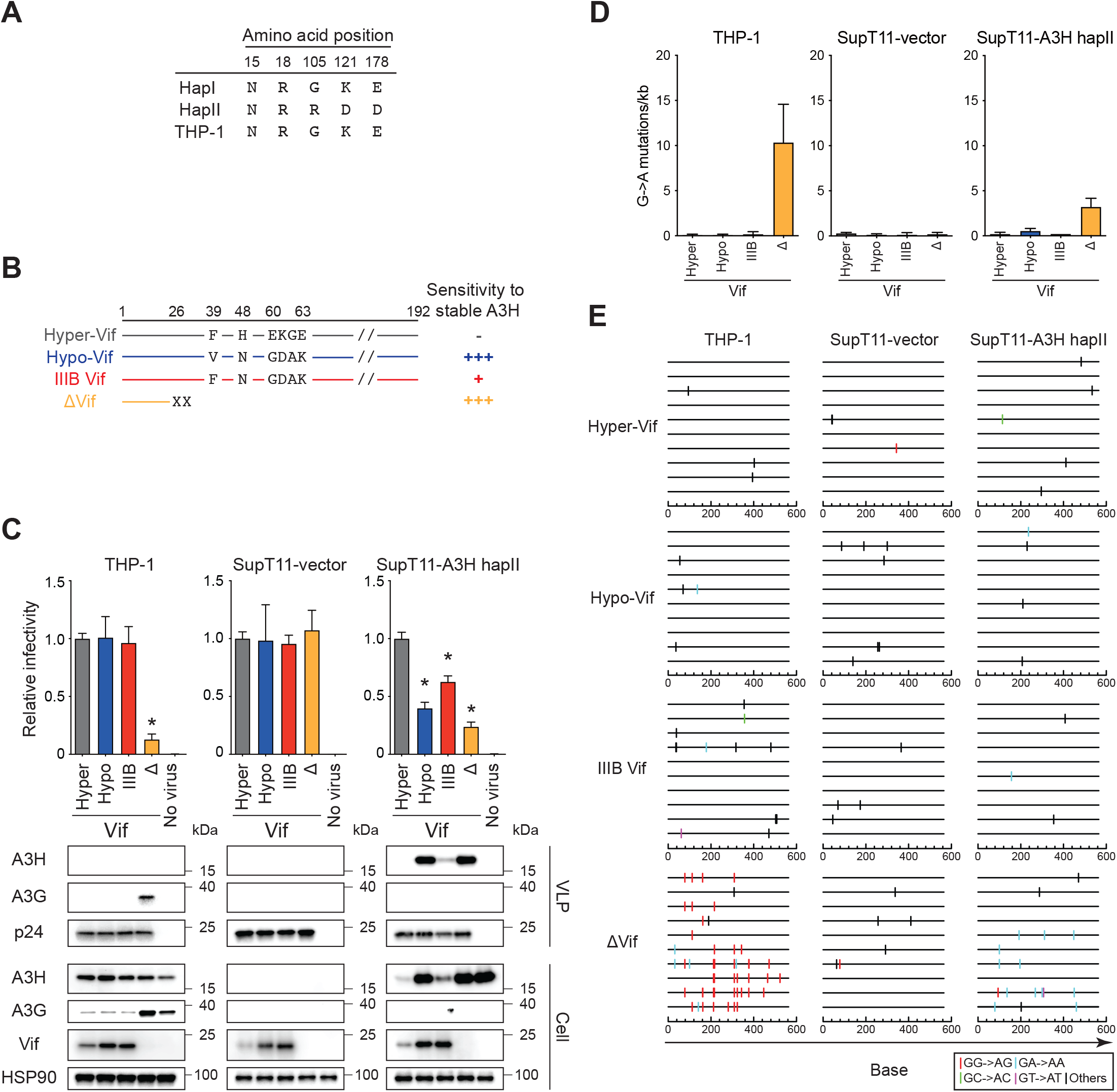
**Endogenous A3H does not inhibit HIV-1 in THP-1 cells.** (**A**) *A3H* haplotypes in THP-1 cells. The indicated positions are key amino acid residues that determine the expression of unstable (hapI) or stable (hapII) A3H protein. (**B**) Schematic of the susceptibility of Vif mutants to stable A3H haplotypes. Key amino acid residues that determine the susceptibility of HIV-1 IIIB Vif to restriction by stable A3H haplotypes. -, full resistance; +, partial resistance; +++, sensitivity. (**C**) Representative infectivity of hyper- and hypo-functional Vif HIV-1 mutants. Top panels show the infectivity of hyper-Vif, hypo-Vif, and IIIB Vif, and Vif-deficient HIV-1 mutants produced in THP-1 cells compared to the same viruses produced in SupT11 cells with stable expression of the control vector or A3H haplotype II. The amounts of produced viruses used to infect TZM-bl cells was normalized to p24 levels. Each bar shows the average of four independent experiments with the standard deviation (SD). Data are represented as relative infectivity compared to hyper-Vif HIV-1. Statistical significance was determined using the two-sided paired *t* test. *P < 0.05 compared with the infectivity of hyper-Vif HIV-1. The bottom panels are representative Western blots of three independent experiments. The levels of viral and cellular proteins in virus-like particles (VLPs) and whole cell lysates are shown. p24 and HSP90 were used as loading controls. (**D**) G-to-A mutations. Average number of G-to-A mutations in the 564 bp *pol* gene after infection with hyper-Vif, hypo-Vif, IIIB Vif, or Vif deficient HIV-1 produced from THP-1 or SupT11 cells expressing either the vector control or A3H hapII. Each bar depicts the average of three independent experiments with SD. (**E**) G-to-A mutation profile. Dinucleotide sequence contexts of G-to-A mutations in the 564 bp *pol* gene after infection with the indicated viruses produced from indicated cell lines. Each vertical line indicates the location of the dinucleotide sequence contexts described in the legend within the 564 bp amplicon (horizontal line).

The A3H hapI results in expression of an unstable protein that has weak anti-HIV-1 activity (28, 29, 31). However, this protein is enzymatically active and has an HIV-1 restriction phenotype similar to the stable A3H haplotype, A3H hapII, when both proteins are expressed at the same levels (31). In addition, A3H protein expression levels are upregulated during HIV-1 infection (10, 22), and A3H hapI is resistant to Vif-mediated degradation (32). Accordingly, we evaluated whether the expression of A3H hapI is associated with HIV-1 restriction in THP-1 cells. To address this question, we utilized HIV-1 Vif mutants that selectively degrade stable A3H (hyper-functional Vif; hyper-Vif) or lack stable A3H degradation (hypo-functional Vif; hypo-Vif) (**Fig. 1B**). IIIB Vif displays an intermediate phenotype (**Fig. 1B**). Of note, hyper-Vif, hypo-Vif, and IIIB Vif have full neutralization activity against A3D, A3F, and A3G proteins (10). VSV-G pseudotyped HIV-1 Vif mutants were produced from HEK293T cells and infected into SupT11 and THP-1 cells to create virus-producing cells (**see Pseudo-single cycle infectivity assays in Material & Methods**). The produced viruses were then used to measure viral infectivity in TZM-bl cells, evaluate packaging of A3 proteins by western blotting, and analyze the frequency of G-to-A mutations. As shown in **Fig. 1C** (**top panel**), hyper-Vif HIV-1, hypo-Vif HIV-1, and IIIB Vif HIV-1 (IIIB) produced in THP-1 cells had similar viral infectivity. While Vif did not degrade A3H protein in THP-1 cells, it was not packaged into viral particles (**Fig. 1C**, **bottom panel**). Next, to determine whether G-to-A mutations were introduced into proviral DNA, we recovered proviral DNA from SupT11 cells after infection with each HIV-1 mutant produced from THP-1 cells and sequenced the *pol* region of these proviruses. Sequencing data demonstrated that hyper-Vif HIV-1, hypo-Vif HIV-1, and IIIB Vif HIV-1 had minimal G-to-A mutations preferred by A3H protein (GA-to-AA signature motif) in proviral DNA (**Fig. 1D and E**), indicating that endogenous A3H protein expressed in THP-1 cells is not involved in HIV-1 restriction. In contrast, the replication of Vif-null HIV-1 was restricted in THP-1 cells and A3G, the major HIV-1 restrictive A3 protein, was packaged in viral particles, thereby inducing profound G-to-A mutations (10.3 ± 3.5 mutations/kb). Most of mutations were in the GG-to-AG signature motif preferred by A3G (80 ± 10%) in proviral DNA (**Fig. 1C-E**). The susceptibility of Vif mutants to stable A3H protein was confirmed in SupT11 cells stably expressing stable A3H protein (**Fig. 1C-E**). Taken together, these results indicate that A3G and other A3 proteins, except A3H, contribute to HIV-1 restriction in THP-1 cells.

### Development of *A3F*-, *A3F/A3G*-, and *A3A*-to-*A3G*-null THP-1 cells

A3F protein has a restrictive effect on HIV-1 among A3 family members and is a target of Vif, in addition to A3G, in CD4^+^ T cell lines and lymphocytes (7, 33-35). To determine whether A3F protein also reduces HIV-1 infectivity in THP-1 cells, we used CRISPR to create *A3F* and *A3F/A3G* gene knockout cell lines. Two independent subclones of *A3F* and *A3F/A3G-*null THP-1 cells were obtained, as evidenced by the results of genomic DNA sequencing and western blotting (**Fig. S1 and S2**).

A3 proteins include single-and double-domain deaminases, which are phylogenetically classified into three groups: Z1, Z2, and Z3 domains (3, 4) (**Fig. 2A represented in green, yellow, and blue, respectively**). A3A, A3B carboxy-terminal domain (CTD), and A3G CTD proteins are classified as Z1 domains (**Fig. 2A; represented in green**). Of note, exon 4 of the *A3A* gene, exon 7 of the *A3B* gene, and exon 7 of *A3G* gene are highly conserved at the nucleotide level (*A3A* exon 4 and *A3B* exon 7 have 95% identity; *A3A* exon 4 and A3G exon 7 have >99% identity; and *A3B* exon 7 and *A3G* exon 7 have 95% identity, respectively). Interestingly, each of these exons has an identical sequence (5′-GAG TGG GAG GCT GCG GGC CA). We therefore designed a guide RNA (gRNA) homologous to this sequence and attempted to delete the entire 125 kbp interval spanning *A3A* to *A3G* in THP-1 cells (**Fig. 2A; represented in arrows, and S3**). We predicted that successful deletion would cause one of the following three scenarios: 1) fusion of exon 4 of the *A3A* gene with exon 7 of the *A3B* gene (30 kbp deletion); 2) fusion of exon 7 of the *A3B* gene with exon 7 of the *A3G* gene (95 kbp deletion); or 3) fusion of exon 4 of the *A3A* gene *with exon 7* of the *A3G* gene (125 kbp deletion; **Fig. 2A**). To obtain THP-1 cells lacking expression of A3A to A3G protein, a lentiviral vector expressing gRNA against the target sequence was transduced into THP-1 cells. Finally, two independent subclones (THP-1#11-4 and THP-1#11-7) were obtained, with whole genome sequencing (WGS) analysis demonstrating an extensive deletion between *A3A* exon 4 and *A3G* exon 7 at the *A3* gene locus (**Fig. 2B**). In THP-1#11-4, six alleles of the fusion of *A3A* exon 4 with *A3G* exon 7 are observed, and each *A3A/A3G* hybrid exon had six different insertions or deletions (indels) (**Fig. S3**). THP-1#11-7 harbors three alleles of *A3A* exon 4 and *A3G* exon 7 fusions (one may be *A3A* exon 4) with three different deletions (**Fig. S3**). Although more than 20 potential off-target sites with two or three nucleotides mismatched with the designed gRNA were predicted, a significant deletion was only found downstream of the predicted *A3G* pseudogene harboring 2 bp mismatched with the target sequence (**Fig. S4**; **potential target sequence in a yellow box and deletions indicated by green dotted lines**). In comparison to parental THP-1 cells, these subclones had similar growth capacities under normal cell culture conditions. RT-qPCR analyses demonstrated that *A3B* to *A3G* mRNA is not detectable in either clone (**Fig. 2C**). However, *A3A* mRNA expression remained detectable in parental THP-1 cells and the two subclones as the A3A promoter remains intact and potentially functional (**Fig. 2A-C**). *A3A* mRNA expression is known to be upregulated 100–1000-fold in THP-1 cell treated with type I interferon (IFN) (36). To confirm the expression of *A3A* mRNA and protein in THP-1 cells, parental THP-1 cell and the respective subclones were cultured in the presence of type I IFN for 6 hours, and *A3* mRNA and protein expression levels were then analyzed by RT-qPCR and western blotting, respectively. In parental THP-1 cells, *A3A*, *A3B*, *A3F*, and *A3G* mRNA and protein expression levels were increased following IFN treatment (**Fig. 2C and D**). In the THP-1#11-4 subclone, *A3A* mRNA expression is increased following IFN treatment; however, A3A, A3B, A3C, A3F, and A3G proteins are not detectable, even after IFN treatment (**Fig. 2C and D**). Further, A3A to A3G proteins are not detectable in the THP-1#11-7 subclone under normal cell culture conditions (**Fig. 2D**). Interestingly, low levels of a protein with comparable size to A3A are detected in the THP-1#11-7 subclone after IFN treatment (**Fig. 2D**). Sanger sequence analyses indicated that this protein was an A3A and A3G hybrid with a 3-bp deletion (**Fig. S3**). Collectively, these data indicate that the THP-1#11-4 and THP-1#11-7 subclones lack expression of A3A to A3G proteins under normal cell culture conditions and that clone THP-1#11-4 is a clean knockout that fails to express functional versions of any of these proteins.

**Figure 2.**
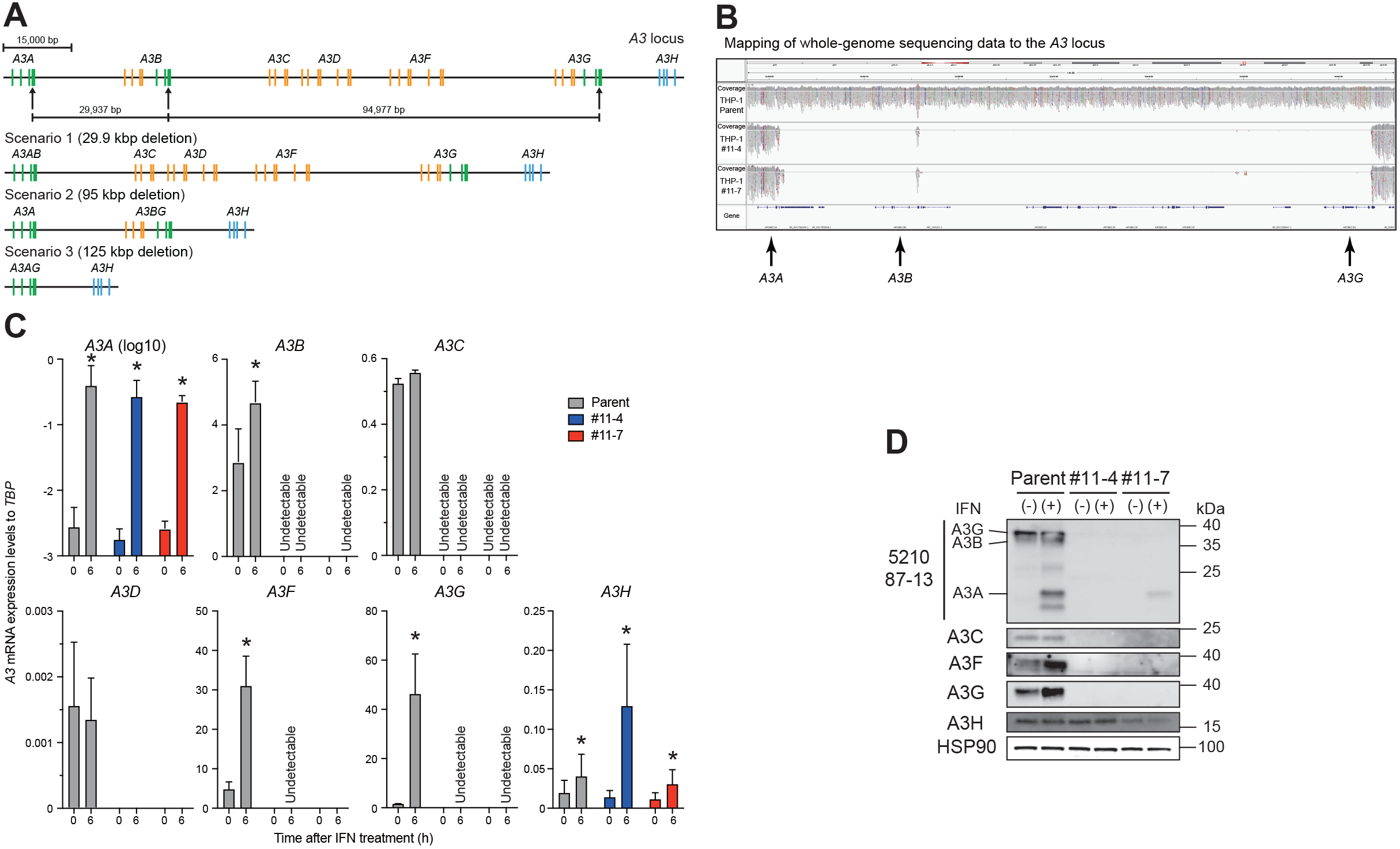
**Disruption of the *A3A* to *A3G* genes in THP-1 cells.** (**A**) Schematic of the A3 *gene* at the *A3* locus. The *A3* family of genes comprises seven members with one or two Z domains (single-or double-domain deaminases) which belong to three phylogenetically distinct groups shown in green, yellow, and blue. Three sites with an identical sequence (5′-GAG TGG GAG GCT GCG GGC CA) in exon 4 of the *A3A* gene, exon 7 of the *A3B* gene, and exon 7 of the *A3G* gene are targeted by gRNA, as indicated by arrows. The three predicted scenarios are shown. Bar represents 15,000 bp. (**B**) Mapping of WGS sequencing data to the *A3* locus. Genomic DNA from parental THP-1, THP-1#11-4, and #11-7 cells were subjected to WGS analysis, with an extensive deletion including the *A3A*–*A3G* genes observed in THP-1#11-4 and #11-7 clones. (**C**) RT-qPCR data. Parental THP-1, THP-1#11-4, and #11-7 cells were treated with 500 units/ml type I IFN. Total RNA was isolated after 6 hours. *A3* mRNA expression levels were quantified by RT-qPCR and are normalized to *TBP* mRNA levels. Each bar represents the average of three independent experiments with SD. Statistical significance was determined using the two-sided paired *t* test. *, P < 0.05 compared to untreated cells. (**D**) Representative Western blots of three independent experiments. Levels of indicated A3 proteins in whole cell lysates from cells treated with or without type I IFN are shown. HSP90 was used as a loading control.

### Disruption of A3A to A3G protein expression fully restores the infectivity of Vif-deficient HIV-1 in THP-1 cells

We next determined whether endogenous A3F protein is degraded by Vif in addition to A3G. HIV-1 Vif mutants with selective A3 neutralization activities were used for pseudo-single cycle infectivity assays as mentioned above. For example, a Vif4A mutant harboring ^14^AKTK^17^ substitutions (^14^DRMR^17^ in IIIB) is susceptible to A3D and A3F activity but resistant to A3G activity (37-39) (**Fig. 3A**). We examined the ability of Vif4A to counteract the activity of A3F as *A3D* mRNA expression level is relatively low in THP-1 cells (26) (**Fig. 2C**). As our group and others have previously shown (26, 37, 38, 40), Vif5A containing five alanine substitutions (^40^YRHHY^44^ to ^40^AAAAA^44^) is sensitive to A3G restriction but not the activity of A3D or A3F (**Fig. 3A**). Vif4A5A is susceptible to A3D, A3F, and A3G (37) (**Fig. 3A**). VSV-G pseudotyped HIV-1 and these Vif mutants were used to infect SupT11 derivatives and engineered *A3F*-null THP-1 cells. First, the susceptibilities of these Vif mutants to A3F and A3G proteins were validated in SupT11 cell lines (**Fig. 3B**). In SupT11-vector cells, Vif-proficient HIV-1 and all Vif mutants had comparable infectivity in TZM-bl cells (**Fig. 3B**). As expected, the infectivity of Vif-deficient HIV-1 and the Vif4A and 4A5A mutants was reduced in SupT11-A3F cells as these mutants are unable to degrade A3F protein, thereby leading to packaging of A3F protein in viral particles (**Fig. 3B**). Further, infection with Vif-deficient HIV-1 or the Vif5A and Vif4A5A mutants resulted in packaging of A3G protein in viral particles from SupT11-A3G cells in addition to reduced infectivity of these Vif mutants (**Fig. 3B**). These results are consistent with previous reports demonstrating the susceptibilities of Vif mutants to A3 proteins (26, 37-40).

**Figure 3.**
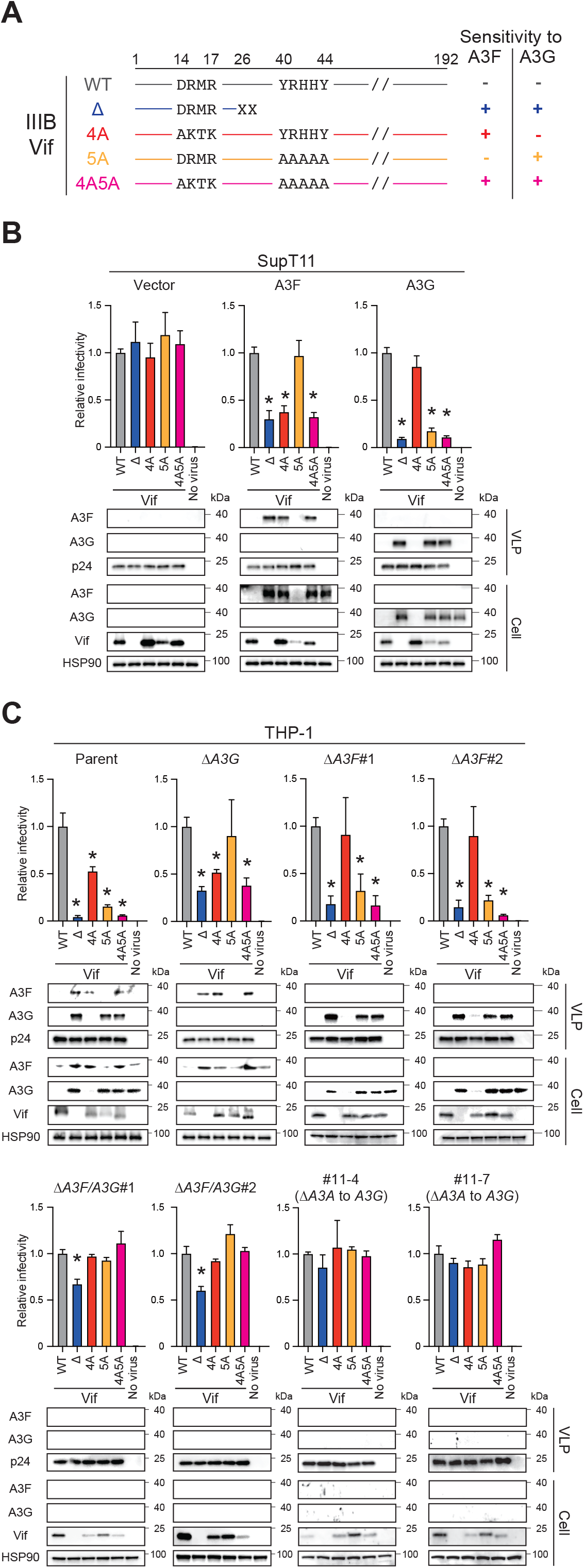
**Pseudo-single cycle infectivity assays for each HIV-1 mutant in *A3*-null THP-1 cells.** (**A**) Schematic of the susceptibility of Vif mutants to A3F and A3G. Key amino acid residues that determine the susceptibility of HIV-1 IIIB Vif to restriction by A3F and A3G. -, resistance; +, sensitivity. (**B**) Representative infectivity of Vif-proficient, Vif-deficient, Vif4A, Vif5A, and Vif4A5A HIV-1 mutants in SupT11 cells stably expressing vector control, A3F, or A3G. Top panels show the infectivity of indicated HIV-1 mutants produced in SupT11 cells stably expressing vector control, A3F, or A3G. The amounts of produced viruses used to infect TZM-bl cells was normalized to p24 levels. Each bar represents the average of four independent experiments with SD. Data are presented as relative infectivity compared to Vif-proficient HIV-1 (WT). Statistical significance was assessed using the two-sided paired *t* test. *P < 0.05 compared to Vif-proficient HIV-1. Bottom panels are representative Western blots of three independent experiments. Levels of indicated viral and cellular proteins in VLPs and whole cell lysates are shown. p24 and HSP90 were used as loading controls. (**C**) Representative infectivity of Vif-proficient, Vif-deficient, Vif4A, Vif5A, and Vif4A5A HIV-1 mutants in *A3*-null THP-1 cells. Top panels show the infectivity of indicated HIV-1 mutants produced in parental or *A3*-null THP-1 cells. The amounts of produced viruses used to infect TZM-bl cells was normalized to p24 levels. Each bar represents the average of four independent experiments with SD. Data are presented as infectivity relative to Vif-proficient HIV-1 (WT). Statistical significance was assessed using the two-sided paired *t* test. *P < 0.05 compared to Vif-proficient HIV-1. Bottom panels are representative Western blots of three independent experiments. Levels of indicated viral and cellular proteins in VLPs and whole cell lysates are shown. p24 and HSP90 were used as loading controls.

Pseudo-single cycle infectivity assays were then performed in parental THP-1, *A3G*-null, and *A3F*-null cells using these Vif mutants. Vif-proficient HIV-1 degraded A3F and A3G proteins in THP-1 cells, and lower amounts of these A3 proteins were packaged into viral particles (**Fig. 3C**; **THP-1 parent**). In contrast, Vif-deficient HIV-1 was unable to degrade A3F and A3G proteins, thereby leading to reduced viral infectivity compared to Vif-proficient HIV-1 (**Fig. 3C**; **THP-1 parent**). The infectivity of A3F-susceptible Vif mutants, Vif4A and Vif4A5A, was lower than that of Vif-proficient HIV-1, indicating that endogenous A3F protein contributes to Vif-deficient HIV-1 restriction in THP-1 cells (**Fig. 3C**; **THP-1 parent**). This finding was supported by results in *A3G*-null THP-1 cells where Vif4A mutants are restricted, as observed in parental THP-1 cells (**Fig. 3C**; **THP-1 `Δ`*A3G).*** The involvement of endogenous A3G protein in HIV-1 restriction was confirmed in *A3G*-null THP-1 cells, as reported (26) (**Fig. 3C**; **THP-1 `Δ`*A3G***). To determine endogenous A3F protein contributes to HIV-1 restriction in THP-1 cells, pseudo-single cycle infectivity assays were performed according to the methods described above in two independent *A3F*-null THP-1 clones (**Fig. S1**). Vif-deficient HIV-1 and the Vif5A and Vif4A5A mutants had reduced infectivity in *A3F*-null subclones due to the inhibitory effect of A3G (**Fig. 3C**; **THP-1 `Δ`*A3F#*1 and #2**). However, the infectivity of the Vif4A mutant was restored to near wildtype levels following disruption of *A3F* expression in THP-1 cells. These data demonstrate that endogenous A3F protein contributes to Vif-deficient HIV-1 restriction in THP-1 cells, and that Vif degrades A3F and thereby prevents packaging and restriction upon target cell infection.

A3F and A3G proteins are involved in Vif-deficient HIV-1 restriction in THP-1 cells and are degraded by Vif (26) (**Fig. 3C**). However, it is unclear whether *only* these A3 proteins are associated with Vif-deficient HIV-1 restriction in THP-1 cells. To address this issue, we performed pseudo-single cycle infectivity assays in *A3F*/*A3G*-null THP-1 cells using separation-of-function Vif mutants. Although Vif-deficient HIV-1 had greater infectivity defects in parental, *A3G*-null, and *A3F*-null THP-1 cells compared to wildtype HIV-1 (parent: <10% infectivity, Δ*A3G*: 30 to 40% infectivity, and Δ*A3F*: 20% infectivity, respectively), the infectivity of Vif-deficient HIV-1 was 30% lower in *A3F*/*A3G*-null THP-1 cells (**Fig. 3C**; **THP-1 parent, `Δ`*A3G*, `Δ`*A3F*#1 and #2, and `Δ`*A3F/A3G*#1 and #2).** On the other hand, the Vif4A, Vif5A, and Vif4A5A mutants had similar infectivity to wildtype HIV-1 in *A3F*/*A3G*-null THP-1 cells (**Fig. 3C; THP-1 `Δ`*A3F/A3G*#1 and #2**). These data indicate that other A3 proteins, in addition to A3F and A3G, contribute to Vif-deficient HIV-1 restriction in THP-1 cells or that Vif disrupts an additional essential target during viral replication in THP-1 cells.

The universally recognized primary target of Vif is the A3 family of proteins (2, 3, 17, 18). However, Vif-mediated A3 degradation may mask an additional A3-independent Vif function required for viral replication. To address this issue, we constructed two independent *A3A*-to-*A3G*-null THP-1 clones (**Fig. 2**) and characterized HIV-1 infection using pseudo-single cycle infectivity assays with Vif mutants. As mentioned above, the disruption of A3F and A3G protein expression results in Vif-deficient HIV-1 having 70% of wildtype HIV-1 infectivity in THP-1 cells (**Fig. 3C**; **THP-1`Δ`*A3F/A3G*#1 and #2).** Remarkably, Vif-deficient HIV-1 and the other Vif mutants have comparable infectivity to Vif-proficient HIV-1 lacking expression of A3A-to-A3G in THP-1 cells (**Fig. 3C**; **THP-1#11-4 and #11-7**). These results indicate that A3 degradation is the only function of Vif required for viral replication in THP-1 cells.

### A3 proteins restrict HIV-1 replication via deaminase-dependent and deaminase-independent mechanisms in THP-1 cells

Our previous results indicated that A3G protein is the primary source of A3 mutagenesis in THP-1 cells (26). To further investigate the G-to-A mutation spectra in each *A3*-null THP-1 subclone, the *pol* region was cloned and sequenced from the proviruses used in the aforementioned infectivity assays. As expected, GG-to-AG mutations are observed in the proviral DNA of Vif mutants lacking A3G neutralization activity (Vif-deficient HIV-1 and Vif5A and Vif4A5A mutants) produced from SupT11-A3G cells (**Fig. 4A-B**; **SupT11-A3G**). Consistent with a previous report (26), THP-1 expresses A3G protein capable of mutating A3G-susceptible Vif mutants, including Vif-deficient HIV-1 and Vif5A and Vif4A5A mutants, as seen in parental THP-1 cells. These GG-to-AG mutations are not observed in *A3G*-null THP-1 cells (**Fig. 4A-B**; **THP-1 parent and `Δ`*A3G***). Similarly, GG-to-AG mutations preferred by A3G were seen in the proviruses of the A3G-susceptible Vif mutants produced from two independent *A3F*-null THP-1 cells, with disruption of A3G nearly completely eliminating these mutations in THP-1 cells (**Fig. 4A and B**; **THP-1`Δ`*A3F*#1 and #2, `Δ`*A3F/A3G*#1 and #2, #11-4, and #11-7**). These data indicate that A3G protein is the primary source of G-to-A mutations in HIV-1 proviruses produced by THP-1 cells.

**Figure 4.**
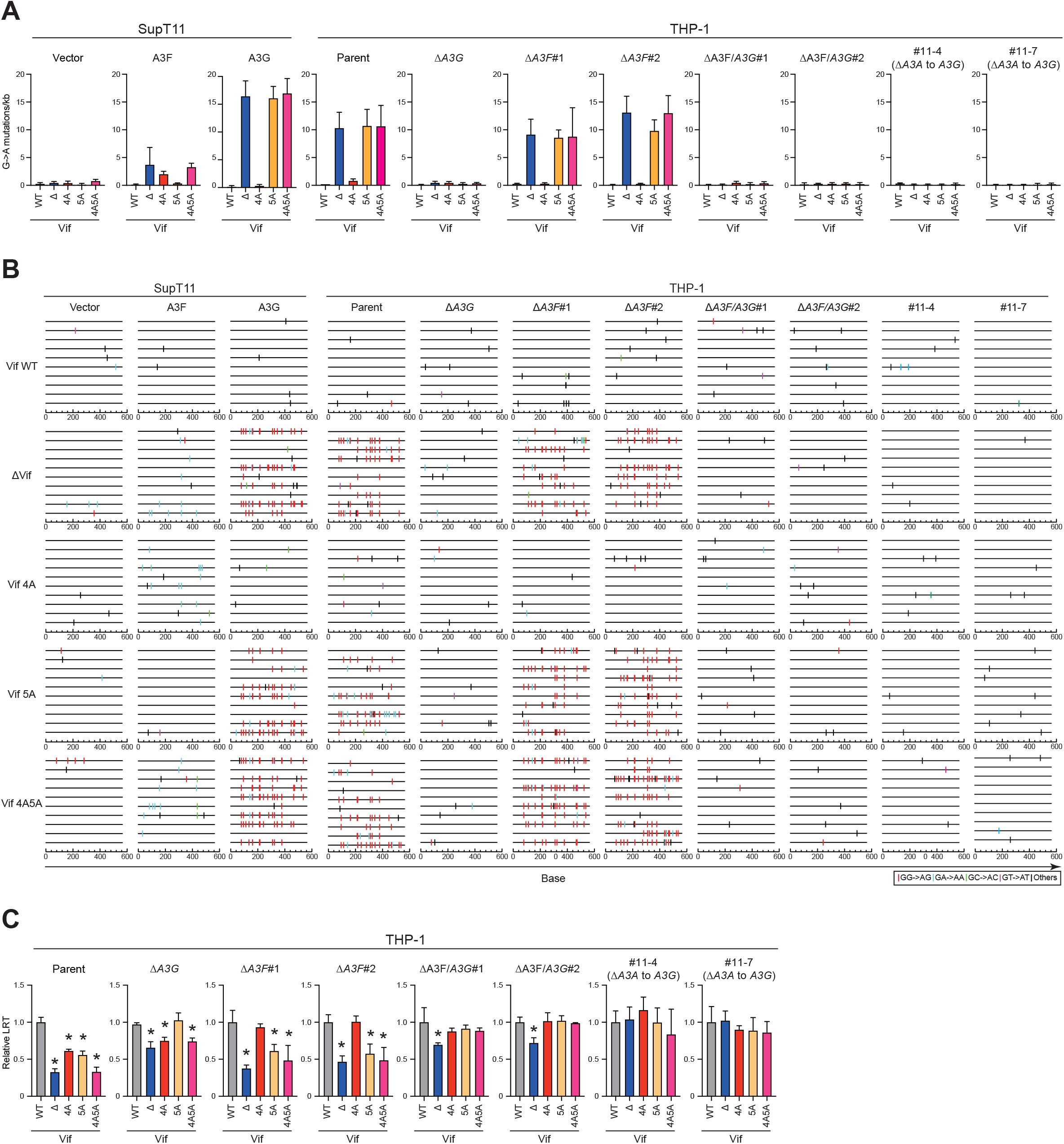
**A3 proteins inhibit Vif-deficient HIV-1 by both deaminase-dependent and independent mechanisms in THP-1 cells.** (**A**)G-to-A mutations. Average number of G-to-A mutations in the 564 bp *pol* gene after infection with hyper-Vif, hypo-Vif, IIIB Vif, or Vif-deficient HIV-1 produced from THP-1 or SupT11 expressing either vector control or A3H hapII. Each bar depicts the average of three independent experiments with SD. (**B**) G-to-A mutation profile. Dinucleotide sequence contexts of G-to-A mutations in the 564 bp *pol* gene after infection with the indicated viruses produced from indicated cell lines. Each vertical line indicates the location of the dinucleotide sequence contexts described in the legend within the 564 bp amplicon (horizontal line). (**C**) Representative LRT quantification data for Vif-proficient, Vif-deficient, Vif4A, Vif5A, and Vif4A5A HIV-1 mutants in each *A3*-null THP-1 subclone. Data show LRT products of the indicated HIV-1 mutants produced in parental or indicated *A3*-null THP-1 cells. The amount of produced viruses used to infect SupT11 cells was normalized to p24 levels. LRT products were measured by qPCR. Each bar represents the average of four independent experiments with SD. LRT products were normalized to the quantity of the *CCR5* gene relative to Vif-proficient HIV-1 (WT). Statistical significance was assessed using the two-sided paired *t* test. *P < 0.05 compared to Vif-proficient HIV-1 LRT products.

Although the Vif mutants lacking A3F neutralization activity (Vif-deficient HIV-1 and Vif4A and Vif4A5A mutants) produced from SupT11-A3F cells have a relatively low number of G-to-A mutations, the observed G-to-A mutations are predominantly within the GA-to-AA sequence motif preferred by A3F (**Fig. 4A-B**; **SupT11-A3F**). However, A3F-preferred GA-to-AA mutations are not observed in proviruses of A3F-susceptible Vif mutants produced from parental or *A3G*-null THP-1 cells, in support of prior observations (26) (**Fig. 4A-B**; **THP-1 parent and `Δ`*A3G***). In addition, fewer GA-to-AA mutations are observed in THP-1 cells, even after disruption of A3F protein expression (**Fig. 4A-B**; **THP-1`Δ`*A3F*#1 and #2, `Δ`*A3F/A3G*#1 and #2, #11-4, and #11-7**). Accordingly, these results combine to indicate that A3F protein in THP-1 cells is involved in Vif-deficient HIV-1 restriction via a deaminase-independent mechanism.

A3F protein has been shown to inhibit the accumulation of reverse transcription (RT) products (14). To investigate a potential effect on RT, SupT11 cells were infected with viruses from the pseudo-single cycle infectivity assays described above, and late RT (LRT) products were examined by quantitative PCR (qPCR). As expected, all Vif mutants were decreased in LRT products in comparison to wildtype virus when these mutants were produced in parental THP-1 cells and used to infect SupT11 cells (**Fig. 4C**; **THP-1 parent**). LRT products of Vif5A and Vif4A mutants were restored to levels comparable to Vif-proficient HIV-1 following the disruption of A3G or A3F protein expression in THP-1 cells (**Fig. 4C**; **THP-1 `Δ`*A3G,* and `Δ`*A3F*#1 and #2**), indicating that both A3G and A3F proteins inhibit HIV-1 via a deaminase-independent mechanism. However, double knockout of A3G and A3F in THP-1 cells did not increase the LRT products of Vif-deficient HIV-1 compared to those of Vif-proficient virus (**Fig. 4C**; **THP-1 `Δ`*A3F/A3G*#1 and #2**), indicating other A3 proteins, in addition to A3F and A3G, may contribute to the restriction of HIV-1 in THP-1 cells via a deaminase-independent mechanism or that a separate protein targeted by Vif blocks the accumulation of RT products. To test this hypothesis, we measured LRT products by infecting SupT11 cells with HIV-1 Vif mutants produced in *A3A*-to-*A3G*-null clones. Consistent with the results of the pseudo-single cycle infectivity assays (**Fig. 3C**), Vif-deficient HIV-1 and other Vif mutants had comparable levels of LRT products to Vif-proficient HIV-1 lacking expression of A3A to A3G protein in THP-1 cells (**Fig. 4C**; **THP-1#11-4 and #11-7**). These data indicate that Vif-mediated A3 degradation is required for viral replication in THP-1 to counteract deaminase-dependent and - independent HIV-1 restriction by A3 proteins.

### Transmitted/founder (TF) HIV-1 Vif also only targets A3 family proteins to enable virus replication in THP-1 cells

We finally examined whether the A3-dependent function of Vif was present in TF viruses. To address this issue, Vif-proficient and deficient versions of the CH58 TF virus were produced from parental THP-1 and *A3A*-to-*A3G*-null cells, with viral infectivity measured in TZM-bl cells (**Fig. 5**). Similar to the results observed with IIIB viruses, Vif-deficient CH58 virus was restricted in parental THP-1 cells; however, this restriction is completely abolished by disruption of the *A3A* to *A3G* genes (**Fig. 5**). These data indicate that TF viruses also utilize a primarily A3-dependent function of Vif during replication in THP-1 cells.

**Figure 5.**
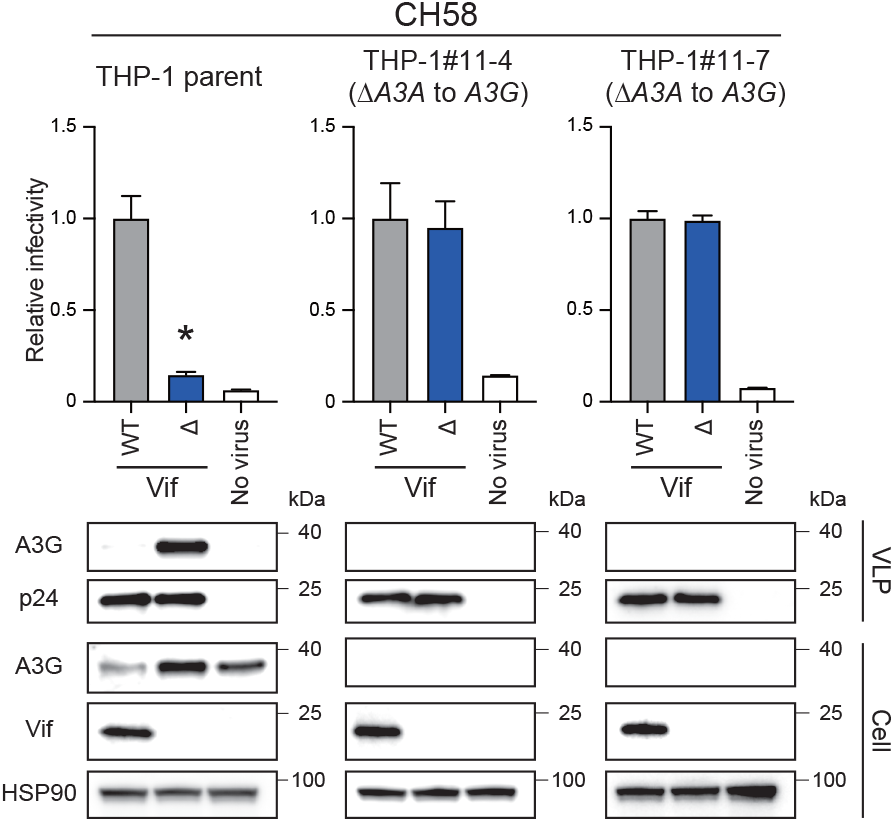
**Pseudo-single cycle infectivity assays of TF virus molecular clone in *A3A-*to*-A3G*-null THP-1 cells.** Infectivity of Vif-proficient and Vif-deficient CH58 viruses. Top panels show the infectivity of Vif-proficient and Vif-deficient HIV-1 produced in parental THP-1, THP-1#11-4, or THP-1#11-7 cells. The amounts of produced viruses used to infect TZM-bl cells was normalized to p24 levels. Each bar represents the average of four independent experiments with SD. Data are represented as relative to Vif-proficient HIV-1 (WT). Statistical significance was assessed using the two-sided paired *t* test. *P < 0.05 compared to Vif-proficient HIV-1. The bottom panels are representative Western blots of three independent experiments. The levels of indicated viral and cellular proteins in VLPs and whole cell lysates are shown. p24 and HSP90 were used as loading controls.

## Discussion

Vif-mediated A3 degradation is critical for HIV-1 replication in CD4^+^ T lymphocytes and myeloid cells (2, 3, 17, 18). In CD4^+^ T lymphocytes, at least A3D, A3F, A3G, and A3H (only stable haplotypes) are involved in Vif-deficient HIV-1 restriction, and Vif is required to degrade A3 enzymes and allow efficient viral replication (2, 3, 17, 18). However, the degradation of A3 enzymes by Vif during HIV-1 replication in myeloid lineage cells has yet to be fully elucidated. We previously reported that A3G protein contributes to Vif-deficient HIV-1 restriction in a deaminase-dependent manner in THP-1 cells (26). Herein, we demonstrate that A3F protein also inhibits Vif-deficient HIV-1 in a largely deaminase-independent manner and that Vif avoids this HIV-1 restriction mechanism by degrading A3F protein (**Fig. 3-4**). Importantly, the results of pseudo-single cycle infectivity assays demonstrate that the disruption of A3A to A3G protein confers comparable infectivity to wildtype HIV-1 in a Vif-deficient lab-adapted virus (IIIB) and TF virus (CH58) (**Fig. 3-5**). These results indicate that Vif-mediated A3 degradation is the primary function of Vif during HIV-1 replication in THP-1 cells.

Our results demonstrate that A3F and A3G but not A3H proteins restrict Vif-deficient HIV-1 via deaminase-dependent and -independent mechanisms in THP-1 cells (**Fig. 1, 3** and **4**). In addition to A3F and A3G proteins, our findings indicate that at least one additional A3 protein is involved in Vif-deficient HIV-1 restriction via a deaminase-independent mechanism (**Fig. 3-4**). Accordingly, the remaining four A3 proteins (A3A, A3B, A3C, and A3D) may contribute to Vif-deficient HIV-1 restriction in a deaminase-independent manner in THP-1 cells (**Fig. 4**). However, A3A and A3B are highly unlikely to contribute in this manner as *A3A* mRNA and protein expression levels are very low or undetectable in THP-1 cells without IFN treatment (**Fig. 2C-D**). Further, both A3A and A3B are resistant to degradation by HIV-1 Vif (7, 34, 41-43). It is therefore plausible that A3C and A3D proteins contribute to Vif-deficient HIV-1 restriction in THP-1 cells. An A3C-isoleucine 188 variant is reportedly more active against HIV-1 than a serine 188 variant (44, 45). To ask which A3C variant is expressed by THP-1 cells, we determined the *A3C* genotypes of THP-1 cells using cDNA sequencing. These results demonstrated that the amino acid residue of A3C at position 188 is serine. This result indicates that A3C has a modest effect on Vif-deficient HIV-1 restriction via a deaminase-independent mechanism in THP-1 cells, consistent with prior studies (45). Similarly, the results of previous studies indicate that A3D has a weak effect on Vif-deficient HIV-1 restriction in HEK293, SupT11, and CEM2n cells (7, 8, 37, 46, 47). Nevertheless, the fact that Vif-deficient HIV-1 has 20% lower infectivity indicates that a synergistic mechanism may enhance the effect of A3 proteins on HIV-1 infectivity (48, 49). Further studies are required to fully elucidate the mechanisms underlying the effect of A3 proteins on HIV-1 infectivity.

Similar to CD4^+^ T lymphocytes, HIV-1 can also target myeloid cells such as monocytes and macrophages, and these infections are associated with viral dissemination, persistence, and latency (50, 51). Accordingly, it is important to understand the role of restriction factors, including A3 proteins, in myeloid cells. In monocytes, *A3A* mRNA levels are 10–1000 times higher than other *A3* mRNA expression levels, and *A3A* mRNA expression is reduced by 10–100-fold after differentiation into monocyte-derived macrophages (MDMs) (52-54). In contrast, *A3G* mRNA expression levels are reduced approximately 10-fold lower after differentiation of monocytes into MDMs (52, 53). *A3F* mRNA expression levels are less variable during the differentiation of monocytes into MDMs (52). Interestingly, suppression of A3A and A3G protein levels by siRNA reportedly leads to a 4–5-fold increase in p24 production by HIV-1-infected monocytes (53). As MDMs are generally more sensitive to HIV-1 infection than monocytes, it is highly likely that A3A and A3G contribute to the susceptibility of MDMs to HIV-1 infection. However, as previous studies have reported that A3A is less active against HIV-1 in HEK293T and SupT11 cell lines (7, 34, 55), further studies are required to determine the contribution of A3A to HIV-1 restriction in monocytes.

In addition to A3A and A3G, A3F and A3H may be involved in HIV-1 restriction in monocytes. Although *A3F* mRNA expression levels are essentially unchanged during differentiation from monocytes into MDMs (53), *A3F* mRNA expression levels are comparable to *A3G* mRNA expression levels (53, 54), indicating that A3F protein likely contributes to HIV-1 restriction in monocytes. It is possible that only stable A3H haplotypes and A3C-I188 are associated with HIV-1 restriction in monocytes. According to previous observations in HEK293, SupT11, and CEM2n cells (7, 8, 37, 46, 47), A3D may modestly contribute to HIV-1 restriction in monocytes. As *A3B* mRNA expression levels are relatively low, it is unlikely that this A3B inhibits HIV-1 in monocytes. However, the contribution of A3 proteins other than A3A and A3G to HIV-1 suppression in monocytes remains unclear, and the antiviral activities of these A3 proteins warrant further investigation.

In MDMs, A3A appears to be associated with anti-HIV-1 activity as increasing HIV-1 infectivity has been reported following siRNA knockdown of *A3A* (53, 54). In addition, HIV-1 replication assays in MDMs using HIV-1 Vif4A and Vif5A mutants demonstrated that the replication kinetics of both mutants were slower than that of the Vif-proficient HIV-1, indicating that A3D, A3F, and A3G contribute to HIV-1 restriction in MDMs (39). However, the effects of A3D and A3F on HIV-1 replication are donor-dependent, likely due to their respective expression levels (39). As the antiviral activity of A3B, A3C, and A3H proteins has not been reported in MDMs, further studies are required to address these issues.

Vif is required for HIV-1 replication in CD4^+^ T lymphocytes and macrophages (2, 3, 17, 18). In the absence of Vif, HIV-1 is attacked by A3 proteins in CD4^+^ T lymphocytes, macrophages, monocytes, dendritic cells, and CD4+ T cell lines, and massive G-to-A mutations accumulate in HIV-1 proviral DNA (7, 8, 10, 15, 23, 26, 39, 56, 57). HIV-1 Vif recruits A3 proteins into an E3 ubiquitin ligase complex, thereby avoiding the antiviral activity of these proteins by promoting their degradation through a proteasome-mediated pathway (2, 3, 17, 18). The primary function of Vif has long been posited to be the suppression of the antiviral activity of A3 proteins. On the other hand, Vif causes G2/M cell cycle arrest (58-60). As the amino acid residues of Vif responsible for G2/M cell cycle arrest do not completely match with the amino acid residues required for Vif-mediated A3 degradation, these functions of Vif may be independent of each other (61-63). In 2016, a functional proteomic analysis identified the PPP2R5 family of proteins, which function as regulators of protein phosphatase 2A (PP2A), as novel targets of Vif (25). Subsequently, Salamango *et al*. revealed that Vif induces G2/M arrest by degrading PPP2R5 proteins (60). Vif-induced G2/M arrest has been observed in many cell types, including HEK293T, SupT11, CEM-SS, and THP-1 cells and CD4^+^ T lymphocytes (25, 61, 63). However, Vif-mediated G2/M arrest is not required for HIV-1 replication, supporting our findings that A3 family proteins are the sole essential substrate of Vif during viral replication in THP-1 cells under normal cell culture conditions (**Fig. 3-5**). It has recently been reported that fragile X mental retardation 1 (FMR1) and diphthamide biosynthesis 7 (DPH7) are degraded by Vif in CD4^+^ T lymphocytes (24). Further studies are required to determine whether a substrate of Vif other than A3 proteins is required for HIV-1 replication in vivo.

In summary, the findings of the present study demonstrate that the primary target of Vif is the A3 family of proteins during HIV-1 replication in THP-1 cells. Whether this observation is applicable to primary CD4^+^ T lymphocytes and myeloid cells, such as monocytes and macrophages, is important for the development of antiviral therapies targeting the A3-Vif axis. Such studies may contribute to a functional cure for HIV-1 by manipulating A3 mutagenesis.

## Material & Methods

### Cell lines and culture conditions

HEK293T (CRL-3216) was obtained from American Type Culture Collection. TZM-bl (#8129) (64) was obtained from the NIH AIDS Reagent Program (NARP). The creation and characterization of the permissive T cell line SupT11 and the SupT11 single clones stably expressing untagged A3 (SupT11-vector, -A3F, -A3G and -A3H hapII high) have been reported (10, 33). CEM-GXR (CEM-GFP expressing CCR5) was provided by Dr. Todd Allen (Harvard University, USA) (65). THP-1 was provided by Dr. Andrea Cimarelli (INSERM, France) (53). The generation and characterization of THP-1 Δ*A3G*#1 have been reported (26). Adherent cells were cultured in DMEM (Wako, Cat# 044-29765) supplemented with 10% fetal bovine serum (FBS) (NICHIREI, Cat#175012) and 1% penicillin/streptomycin (P/S) (Wako, Cat# 168-23191). Suspension cells were maintained in RPMI (Thermo Fisher Scientific, Cat# C11875500BT) with 10% FBS and 1% P/S.

### Genotyping of *A3C* and *A3H* genes

Total RNA was isolated from THP-1 by RNA Premium Kit (NIPPON Genetics, Cat# FG-81250). Then, cDNA was synthesized by Transcriptor Reverse Transcriptase (Roche, Cat# 03531287001) and used to amplify *A3C* or *A3H* gene with the following primers [*A3C* outer primers: (5’-GCG CTT CAG AAA AGA GTG GG) and (5’-GGA GAC AGA CCA TGA GGC). *A3C* inner primers: (5’-ACA TGA ATC CAC AGA TCA GAA A) and (5’-CCC CTC ACT GGA GAC TCT CC). *A3H* outer primers: (5’-CCA GAA GCA CAG ATC AGA AAC ACG AT) and (5’-GAC CAG CAG GCT ATG AGG CAA). *A3H* inner primers: (5’-TGT TAA CAG CCG AAA CAT TCC) and (5’-TCT TGA GTT GCT TCT TGA TAA T)]. The amplified fragments were cloned into the pJET cloning vector (Thermo Fisher Scientific, Cat# K1231). At least 10 independent clones were subjected to Sanger sequencing (AZENTA) and sequence data were analyzed by Sequencher v5.4.6 (Gene Codes Corporation).

### Construction of pLentiCRISPR-Blast

The pLentiCRISPR1000 system was previously described (66). pLentiCRISPR1000-Blast was generated by restriction digest with BmtI and MluI to excise the P2A-puromycin cassette. An oligo containing a P2A-blasticidin cassette was purchased from IDT (5’-AGC GGA GCT ACT AAC TTC AGC CTG CTG AAG CAG GCT GGC GAC GTG GAG GAG AAC CCT GGA CCT ACC GGT ATG GCC AAG CCA CTG TCC CAA GAA GAG TCA ACT CTG ATC GAG AGG GCC ACT GCA ACC ATT AAT AGC ATT CCC ATC TCT GAA GAC TAT AGC GTA GCT AGT GCC GCA CTC AGC TCT GAT GGA CGC ATA TTC ACC GGC GTT AAT GTC TAC CAC TTC ACC GGC GGA CCC TGC GCC GAA CTG GTC GTG CTG GGG ACC GCA GCC GCC GCG GCT GCC GGG AAT TTG ACG TGC ATT GTT GCA ATA GGC AAC GAG AAT AGG GGC ATC CTG TCA CCT TGC GGC CGG TGT CGG CAA GTG CTG CTG GAC CTG CAC CCC GGC ATC AAG GCC ATA GTC AAG GAT AGT GAT GGC CAG CCG ACC GCC GTT GGG ATT CGA GAA CTT CTG CCT TCT GGG TAC GTC TGG GAA GGC TAG) and amplified with the primers (5’-CAA GAC TAG TGG AAG CGG AGC TAC TAA CTT CAG CCT GCT GAA GCA GGC TGG CGA CGT GGA GGA and 5’-NNN NAC GCG TCT AGC CTT CCC AGA CGT ACC C) using high-fidelity Phusion polymerase (NEB, Cat# M0530S). The PCR fragment was digested with BmtI and MluI, and ligated into the cut pLentiCRISPR1000, producing pLentiCRISPR1000-Blast.

### Creation of THP-1 cells disrupting *A3* genes

An *A3F* specific guide for *exon 3* was designed (**Fig. S1A and S2A**) and evaluated manually for specificity to the *A3F* target sequence via an alignment with the most related members of the *A3* family as described previously (26). Oligos with ends compatible with the Esp3I sites in pLentiCRISPR1000-Blast were purchased from IDT [Δ*A3F* gRNA: (5’-CAC CGG TAG TAG TAG AGG CGG GCG G) and (5’-CCA TCA TCA TCT CCG CCC GCC CAA G)]. The targeting construct was generated by annealing oligos and cloned by Golden Gate ligation into pLentiCRISPR1000-Blast. A guide with a common sequence among *A3A exon 4*, *A3B exon 7* and *A3G exon 7* was designed (**Fig. 2A**) and oligos with ends compatible with the Esp3I sites in pLentiCRISPR1000 (66) were purchased from IDT [PanZ1 gRNA: (5’-CAC CGT GGC CCG CAG CCT CCC ACT C) and (5’-GAA CGA GTG GGA GGC TGC GGG CCA C)]. The targeting construct was generated by annealing oligos and cloned by Golden Gate ligation into pLentiCRISPR1000 (66). All constructs were confirmed by Sanger sequencing (AZENTA) and sequence data were analyzed by Sequencher v5.4.6 (Gene Codes Corporation).

For transduction, VSV-G pseudotyped virus was generated by transfecting 2.5 μg of the pLentiCRISPR1000 or pLentiCRISPR1000-Blast targeting construct along with 1.67 μg of pΔ-NRF (HIV-1 *gag*, *pol*, *rev*, *tat* genes) (67) and 0.83 μg of pMD.G (VSV-G) expression vectors using TransIT-LT1 (Takara, Cat# MIR2306) into 293T cells. At 48 hours post-transfection, viral supernatants were harvested, filtered with 0.45 µm filters (Merck, Cat# SLHVR33RB), and concentrated by centrifugation (26,200 × *g*, 4°C, 2 hours). Then, viral pellets were resuspended in 10% FBS/RPMI and incubated with cells for 48 hours. Forty-eight hours later, cells were placed under drug selection in 10% FBS/RPMI containing 1 µg/ml puromycin (InvivoGen, Cat# ant-pr) or 6 ng/ml blasticidin (InvivoGen, Cat# ant-bl). Single-cell clones were isolated by the limiting dilution of the drug-resistant cell pool and expanded. The expression levels of A3F protein in THP-1 Δ*A3F*#1 and #2, and THP-1Δ*A3F/A3G*#1 and #2 cells were confirmed by immunoblots (see Western blots). To confirm indels in the *A3F* target sequence of the selected clones, genomic DNA was isolated by DNeasy Blood & Tissue Kits (Qiagen, Cat# 69504) and amplified with Choice-Taq DNA polymerase (Denville Scientific, Cat# CB4050-2) using primers (5’-GCT GAA GTC GCC CTT GAA TAA ACA CGC and 5’-TGT CAG TGC TGG CCC CG). The amplified PCR products were cloned into the pJET cloning vector (Thermo Fisher Scientific, Cat# K1231) and subjected to Sanger sequencing (AZENTA). To confirm indels in the *A3A, A3B and A3G* target sequences of the selected clones (THP-1#11-4 and #11-7), genomic DNA was isolated by DNeasy Blood & Tissue Kits (Qiagen, Cat# 69504) and subjected to whole genome sequencing (WGS) (macrogen). The sequencing data were aligned by Isaac aligner (iSAAC-04.18.11.09). Off-target sites were analyzed by Cas-OFFinder (http://www.rgenome.net/cas-offinder/). For further analysis of indels between *A3A* and *A3G*, genomic DNAs from THP-1#11-4 and #11-7 were amplified using primers (5’-GGG GCT TTC TGA AAG AAT GAG AAC TGG GC and 5’-CAG CTG GAG ATG GTG GTG AAC AGC C). The amplified PCR products were cloned into the pJET cloning vector (Thermo Fisher Scientific, Cat# K1231) and subjected to Sanger sequencing (AZENTA). All sequence data were analyzed by Sequencher v5.4.6 (Gene Codes Corporation). To assess the expression levels of *A3* mRNAs and proteins, THP-1 parent, #11-4, and #11-7 were incubated in 10%FBS/RPMI including 500 units/ml IFN (R & D Systems, Cat# 11200-2) for 6 hours. Then, cells were harvested and subjected to RT-qPCR (see RT-qPCR) (**Fig. 2C**) and Western blot (see Western blot) (**Fig. 2D**).

### Pseudo-single cycle infectivity assays

Vif-proficient and Vif-deficient (X^26^ and X^27^) HIV-1 IIIB C200 proviral expression constructs have been reported (68). HIV-1 IIIB C200 mutants with hyper- (H^48^ and ^60^EKGE^63^) and hypo- (V^39^) functional Vifs have been reported (10). An HIV-1 IIIB C200 Vif 5A mutant (^40^AAAAA^44^) has been described (26). HIV-1 IIIB C200 Vif 4A (14AKTK18) and 4A5A (^14^AKTK^18^ and ^40^AAAAA^44^) mutants were created by digesting pNLCSFV3-4A, and -4A5A proviral DNA construct [(37); kindly provided by Dr. Kei Sato, University of Tokyo, Japan] at SwaI and SalI sites and cloned into pIIIB C200 proviral construct. The proviral expression vector encoding full length TF virus, CH58 (#11856) was obtained from the NARP. The creation of Vif-deficient CH58 mutant has been described previously (69).

HIV-1 single-cycle assays using VSV-G pseudotyped viruses were performed as described previously (23, 26). 293T cells were cotransfected with 2.4 μg of proviral DNA construct and 0.6 μg of VSV-G expression vector using TransIT-LT1 reagent (Takara, Cat# MIR2306) into 293T cells (3 × 10^6^). Forty-eight hours later, supernatants were harvested, filtered (0.45 μm filters, Merck, Cat# SLHVR33RB), and used to titrate on 2.5 × 10^4^ CEM-GXR reporter cells for MOI determinations. GFP+ cells were measured using a FACS Canto II (BD Biosciences) and the data were analyzed using FlowJo software v10.7.1 (BD Biosciences). 1 or 5 × 10^6^ target cells were infected with an MOI of 0.05 (for SupT11 derivatives) or 0.25 (for THP-1 derivatives) and washed with PBS twice at 24 hours post-infection and then incubated for an additional 24 hours. After 24 hours, supernatants were collected and filtered. The resulting viral particles were quantified by p24 ELISA (ZeptoMetrix, Cat# 0801008) and used to infect 1 × 10^4^ TZM-bl cells (1 or 2 ng of p24). At 48 hours postinfection, the infected cells were lysed with a Bright-Glo luciferase assay system (Promega, Cat# E2650) and the intracellular luciferase activity was measured by a Synergy H1 microplate reader (BioTek) or Centro XS3 LB960 microplate luminometer (Berthold Technologies).

### Quantification of LRT products

Viruses were produced by infecting VSV-G pseudotyped virus into THP-1 cells as described above (see HIV-1 infectivity assays) and the resulting viral particles were quantified by p24 ELISA (ZeptoMetrix, Cat# 0801008). The viral supernatants including 20 ng of p24 antigen were used for infection into SupT11 cells. At 12 hours postinfection, cells were harvested and washed with PBS twice. Then, total DNA was isolated by DNeasy Blood & Tissue Kits (Qiagen, Cat# 69504) and treated with RNase A (Qiagen, Cat# 19101) according to the manufacturer’s instruction. Following DpnI digestion, 50 ng of DNA was used to amplify LRT products and *CCR5* gene with the following primers; LRT forward: (5’-CGT CTG TTG TGT GAC TCT GG) and LRT reverse: (5’-TTT TGG CGT ACT CAC CAG TCG). *CCR5* forward: (5’-CCA GAA GAG CTG AGA CAT CCG) and *CCR5* reverse (5’-GCC AAG CAG CTG AGA GGT TAC T). qPCR was performed using Power SYBR Green PCR Master Mix (Thermo Fisher Scientific, Cat# 4367659) and fluorescent signals from resulting PCR products were acquired using a Thermal Cycler Dice Real Time System III (Takara). Finally, each LRT product was represented as values normalized by the quantity of the *CCR5* gene (**Fig. 4C**).

### RT-qPCR

Cells were harvested and washed with PBS twice. Then, total RNA was isolated by RNA Premium Kit (NIPPON Genetics, Cat# FG-81250) and cDNA was synthesized by Transcriptor Reverse Transcriptase (Roche, Cat# 03531287001) with random hexamer. RT-qPCR was performed using Power SYBR Green PCR Master Mix (Thermo Fisher Scientific, Cat# 4367659). Primers for each *A3* mRNA have been reported previously (70, 71). *A3A* forward: (5’-GAG AAG GGA CAA GCA CAT GG) and *A3A* reverse: (5’-TGG ATC CAT CAA GTG TCT GG). *A3B* forward: (5’-GAC CCT TTG GTC CTT CGA C) and *A3B* reverse: (5’-GCA CAG CCC CAG GAG AAG). *A3C* forward: (5’-AGC GCT TCA GAA AAG AGT GG) and *A3C* reverse: (5’-AAG TTT CGT TCC GAT CGT TG). *A3D* forward: (5’-ACC CAA ACG TCA GTC GAA TC) and *A3D* reverse: (5’-CAC ATT TCT GCG TGG TTC TC). *A3F* forward: (5’-CCG TTT GGA CGC AAA GAT) and *A3F* reverse: (5’-CCA GGT GAT CTG GAA ACA CTT). *A3G* forward: (5’-CCG AGG ACC CGA AGG TTA C) and *A3G* reverse: (5’-TCC AAC AGT GCT GAA ATT CG). *A3H* forward: (5’-AGC TGT GGC CAG AAG CAC) and *A3H* reverse: (5’-CGG AAT GTT TCG GCT GTT). *TATA-binding protein* (*TBP*) forward: (5’-CCC ATG ACT CCC ATG ACC) and *TBP* reverse: (5’-TTT ACA ACC AAG ATT CAC TGT GG). Fluorescent signals from resulting PCR products were acquired using a Thermal Cycler Dice Real Time System III (Takara). Finally, each *A3* mRNA expression level was represented as values normalized by *TBP* mRNA expression levels (**Fig. 2C**).

### Hypermutation analyses

Hypermutation analyses were performed as previously described (23, 26, 45). Genomic DNAs containing HIV-1 proviruses were recovered by infecting viruses produced in derivatives of THP-1 or SupT11 cells into SupT11 using DNeasy Blood & Tissue Kits (Qiagen, Cat# 69504). Following DpnI digestion, the viral *pol* region was amplified by nested PCR with outer primers (876 bp) [(5’-TCC ART ATT TRC CAT AAA RAA AAA) and (5’-TTY AGA TTT TTA AAT GGY TYT TGA)] and inner primers (564 bp) [(5’-AAT ATT CCA RTR TAR CAT RAC AAA AAT) and (5’-AAT GGY TYT TGA TAA ATT TGA TAT GT)]. The resulting 564 bp amplicon was subjected to pJET cloning. At least 10 independent clones were Sanger sequenced (AZENTA) for each condition and analyzed by the HIV sequence database (https://www.hiv.lanl.gov/content/sequence/HYPERMUT/hypermut.html). Clones with identical mutations were eliminated.

### Western blot

Western blot for cell and viral lysates were performed as described previously (23, 26, 72). Cells were harvested, washed with PBS twice, and lysed in lysis buffer [25 mM HEPES (pH7.2), 20% glycerol, 125 mM NaCl, 1% Nonidet P40 (NP40) substitute (Nacalai Tesque, Cat# 18558-54)]. After quantification of total protein by protein assay dye (Bio-Rad, Cat# 5000006), lysates were diluted with 2 × SDS sample buffer [100 mM Tris-HCl (pH 6.8), 4% SDS, 12% β-mercaptoethanol, 20% glycerol, 0.05% bromophenol blue] and boiled for 10 minutes. Virions were dissolved in 2 × SDS sample buffer and boiled for 10 minutes after pelleting down using 20% sucrose (26,200 × *g*, 4°C, 2 hours). Then, the quantity of p24 antigen was measured by p24 ELISA (ZeptoMetrix, Cat# 0801008).

Proteins in the cell and viral lysates (5 μg of total protein and 10 ng of p24 antigen) were separated by SDS-PAGE and transferred to PVDF membranes (Millipore, Cat# IPVH00010). Membranes were blocked with 5% milk in PBS containing 0.1% Tween 20 (0.1% PBST) and incubated in 4% milk/0.1% PBST containing primary antibodies: mouse anti-HSP90 (BD Transduction Laboratories, Cat# 610418, 1:5,000); rabbit anti-A3B (5210-87-13, 1:1,000) (73); rabbit anti-A3C (Proteintech, Cat# 10591-1-AP, 1:1,000); rabbit anti-A3F (675, 1:1,000) (74); rabbit anti-A3G (NARP, #10201, 1:2,500); rabbit anti-A3H (Novus Biologicals, NBP1-91682, 1:5,000): mouse anti-Vif (NARP, #6459, 1:2,000); mouse anti-p24 (NARP, #1513, 1:2,000). Subsequently, the membranes were incubated with horseradish peroxidase (HRP)-conjugated secondary antibodies: donkey anti-rabbit IgG-HRP (Jackson ImmunoResearch, 711-035-152; 1:5,000); donkey anti-mouse IgG-HRP (Jackson ImmunoResearch, 715-035-150). SuperSignal West Femto Maximum Sensitivity Substrate (Thermo Fisher Scientific, Cat# 34095) or Super signal atto (Thermo Fisher Scientific, Cat# A38555) was used for HRP detection. Bands were visualized by the Amersham Imager 600 (Amersham).

### Statistical analyses

Statistical significance was performed using a two-sided paired *t* test (Fig. 1C, 2C, 3B, 3C, 4C, **and 5**). GraphPad Prism software v8.4.3 was used for these statistical tests.

## Acknowledgements

We would like to thank Haruyo Hasebe, Kimiko Ichihara, Kazuko Kitazato, Otowa Takahashi, and all Ikeda lab members for technical assistance. We also would like to thank Drs. Todd Allen, Andrea Cimarelli, and Kei Sato for sharing reagents. This study was supported in part by AMED Research Program on Emerging and Re-emerging Infectious Diseases (JP21fk0108574, to Hesham Nasser; JP21fk0108494 to Terumasa Ikeda); AMED Research Program on HIV/AIDS (JP22fk0410055, to Terumasa Ikeda); JSPS KAKENHI Grant-in-Aid for Scientific Research C (22K07103, to Terumasa Ikeda); JSPS KAKENHI Grant-in-Aid for Early-Career Scientists (22K16375, to Hesham Nasser); JSPS Leading Initiative for Excellent Young Researchers (LEADER) (to Terumasa Ikeda); Takeda Science Foundation (to Terumasa Ikeda); Mochida Memorial Foundation for Medical and Pharmaceutical Research (to Terumasa Ikeda); The Naito Foundation (to Terumasa Ikeda); Shin-Nihon Foundation of Advanced Medical Research (to Terumasa Ikeda); Waksman Foundation of Japan (to Terumasa Ikeda); an intramural grant from Kumamoto University COVID-19 Research Projects (AMABIE) (to Terumasa Ikeda); Intercontinental Research and Educational Platform Aiming for Eradication of HIV/AIDS (to Terumasa Ikeda); International Joint Research Project of the Institute of Medical Science, the University of Tokyo (to Terumasa Ikeda); SPP1923 and the Heisenberg Program of the German Research Foundation (SA 2676/3-1; SA 2676/1-2) (to Daniel Sauter), as well as the Canon Foundation Europe (to Daniel Sauter). Contributions from the Harris lab were supported by NIAID R37-AI064046. RSH is an Investigator of the Howard Hughes Medical Institute, a CPRIT Scholar, and the Ewing Halsell President’s Council Distinguished Chair.

The authors declare that they have no competing interests.

---

**Figure S1. Development of *A3F*-null THP-1 cells**.

(**A**) *A3F* exon 3 sequences encompassing the gRNA target site in parental THP-1 and two independent *A3F*-null THP-1 cells. Indels in two alleles for each *A3F*-null THP-1 clone are shown.

(**B**) Representative Western blots of three independent experiments. Levels of A3F and A3G protein in whole cell lysates are shown. HSP90 was used as a loading control.

**Figure S2. Development of *A3F/A3G*-null THP-1 cells.**

(**A**) *A3F* exon 3 sequences encompassing the gRNA target site in parental THP-1 and two independent *A3F/A3G*-null THP-1 cells. Indels in two alleles for each *A3F/A3G*-null THP-1 clone are shown.

(**B**) Representative Western blots of three independent experiments. Levels of A3F and A3G protein in whole cell lysates are shown. HSP90 was used as a loading control.

**Fig. S3 Sequence analysis of flanking region targeted by gRNA in THP-1#11-4 and #11-7.**

(**A**) *A3A* exon 4 and *A3G* exon 7 hybrid sequences encompassing the gRNA target site in THP-1#11-4 cells. Only one nucleotide difference (>99% identity) was observed between *A3A* exon 4 and *A3G* exon 7 and is shown in purple (A3A, cytosine) or green (A3G, adenine). Indels in six alleles of the THP-1#11-4 clone are shown.

(**B**) *A3A* exon 4 and *A3G* exon 7 hybrid sequences encompassing the gRNA target site in THP-1#11-7 cells. Only one nucleotide difference (>99% identity) was observed between *A3A* exon 4 and *A3G* exon 7 and is shown in purple (A3A, cytosine) or green (A3G, adenine). Indels in three alleles of the THP-1#11-7 clone are shown.

**Fig. S4 Deletions around predicted *A3G* pseudogene.**

Mapping of WGS sequencing data to off-target and downstream regions on chromosome 12. Genomic DNA from parental THP-1, THP-1#11-4, and THP-1#11-7 cells were subjected to WGS analysis. The yellow box indicates the off-target sequence in the predicted pseudogene. Several deletions were observed in the regions indicated by green dot boxes in THP-1#11-4 and THP-1#11-7 clones.

